# Potentiation of host defense through sRNA packaged in OMVs of *Xanthomonas oryzae* pv. *oryzicola*

**DOI:** 10.1101/2023.03.10.532040

**Authors:** Yan Wu, Sai Wang, Peihong Wang, Wenhan Nie, Iftikhar Ahmad, Gongyou Chen, Bo Zhu

## Abstract

Bacterial outer membrane vesicles (OMVs)-packaged delivery of noncoding small RNAs (sRNAs) can function as novel mediators of interspecies communication. However, the role of which in the interaction between phytopathogenic bacteria and their host plants is unclear. In this study, we characterized differentially packaged sRNAs in *Xanthomonas oryzae* pv. *oryzicola* (*Xoc*) BLS256 OMVs using RNA-Seq, and we selected the most abundant sRNA Xosr001 for further study based on its essential role in the induction of stomatal immunity in rice. *Xoc* loads Xosr001 into OMVs, which are transferred specifically into the mechanical tissues of rice leaves. We uncovered that OMVs-mediated Xosr001 inhibitors attenuated *OsJMT1* transcripts in vivo and reduced the endogenous MeJA contents in rice leaves. Stomatal conductance was measured to show that ΔXosr001 mutant weakened the ability of stomatal re-opening on rice leaves after spray inoculation. Most notably, the transgenic rice lines OsJMT1-HA-OE exhibited attenuated stomatal immunity and disease susceptibility after ΔXosr001 infection compared with *Xoc* infection. These results define that Xosr001 packaged in *Xoc* OMVs highlights a smart molecular mechanism to activate stomatal immunity in rice.

## Introduction

Bacterial outer membrane vesicles (OMVs) play an essential role for interspecies and interkingdom interactions, which constantly divorced and formed from outermost membrane of bacteria during its life activity [1–3]. OMVs act as nanospherical proteoliposomes with a diameter ranging from 20 to 400 nm, inside them contain protein, lipid, nucleic acid and other ingredient, which deliver functional molecules to host cells to participate in immunization strategies [4, 5]. Interestingly, small RNAs (sRNAs) can be internalized within OMVs, transferred into host cytoplasm, and help to establish interspecies communication [6, 7]. Due to the demonstrated capacity of sRNA-mediated post-transcriptional gene regulation in hosts, sRNAs released from OMVs have been considered as candidate molecules for bacteria-host interaction. Bacterial sRNA exerts multiple biological functions, including environmental adaptation, quorum sensing, bacterial communication and virulence, mainly by regulating the transcription level of target genes through complementary base pairing [8–10].

Several studies have proved that host immune responses triggered by sRNA packaged in OMVs occur during bacterial invasion, and favor both in vitro bacterial growth and host infection [6, 11–13]. For instance, sRNA SsrA is specifically transported into the epithelial cells of squid *Euprymna scolopes* via OMVs and modulate the retinoic-acid inducible gene-I (RIG-I) like receptor signaling pathway in *Vibrio fischeri* [12]. Similarly, sRNA52320 can be packed into *Pseudomonas aeruginosa* OMVs, which transferred into human epithelial airway cells and reduced LPS-induced IL-8 secretion [6].

Though the majority of studies on OMVs have been reported extensively for animal-pathogenic bacteria, yet little is known about the mechanistic role of OMVs for the interaction between phytopathogenic bacteria and host plants. The biological strategy of OMV production used by phytopathogenic bacteria favored either the bacterial life cycle inside host plants or the activation of host immune responses. *Xylella fastidiosa* OMVs fine-tune attachment to surfaces *in planta*, thereby facilitating the binding to the walls of xylem vessels and contributing to virulence [14]. *Pseudomonas syringae* pv. *tomato* DC3000 OMVs elicit plant immune responses which showed that OMV-mediated of plant immunity against the fungal pathogen, as well as against bacterium [15, 16]. Recently, the study reported the immune triggering property of OMVs produced by *Xanthomonas campestris* pv. *campestris* (*Xcc*), which could directly insert into the Arabidopsis plasma membrane (PM) to enhance the lipid order of plant PM and thus strengthen the plant defenses [17]. However, the role of sRNA packaged in OMVs playing in plant-microbe interaction is poorly characterized, especially in phytopathogenic bacteria.

*Xanthomonas oryzae* pv. *oryzicola* (*Xoc*), causing bacterial leaf streak in rice, infects rice leaves through stomata and wounds [18]. Stomatal immunity develops during the early stages of bacterial infection [19]. Phytopathogenic bacteria secrete effectors or phytotoxins to suppress stomatal immunity. For example, *Xoc* type III effector XopC2 activates JA signaling, thereby suppressing stomatal closure to facilitate bacterial entry of rice leaves [20].

In this study, we focused on the mechanism that if sRNAs packaged in OMVs of the gram-negative plant pathogen *Xoc* BLS256 could load into host plant cells and how sRNAs affects plant immune responses. OMVs released from BLS256 were extracted by ultracentrifugation and purified by density gradient centrifugation. RNA-Seq and northern blot were used to identify four candidate sRNAs packaged in OMVs. Here, we provide a first characterization of *Xoc* BLS256 sRNA (Xosr001), the most abundant sRNA within OMVs, which could be transferred from OMVs to host plant cells and located in the epidermis, mechanical tissue and phloem of rice leaves, where it attenuates the transcriptional level of *OsJMT1* and results in reducing the accumulation of MeJA in rice leaves. The construction of OsJMT1-overexpressing and OsJMT1-knockout transgenic lines in rice cv. Nippobare were used to evaluate that Xosr001 impacted *OsJMT1* transcription. In addition, stomatal conductance analysis and spray inoculation assays showed that OMVs lacking Xosr001 greatly restrict bacterial infection at the very early infection stage whereby Xosr001 packed inside OMVs of *Xoc* BLS256 heightened stomatal immunity to potentiate plant defense in rice.

## Materials and methods

### Strains, plasmids and primers

The bacterial strains and plasmids used in this study are described in Table S1. Primers used for the construction of mutant strains, plasmids and DNA templates are provided in Table S2.

### Plant materials and bacterial strains

*Oryza sativa* ssp. *japonica* cv. Nipponbare was used as wild type and for generating transgenic plants. *Escherichia coli* DH5α was cultured in Luria-Bertani (LB) medium at 37°C. *Xoc* BLS256 and derivative strains were grown in nutrient broth (NB) or NB containing 1.5% (w/v) agar (NA) as described previously [21]. YEB medium (5 g/l yeast extract powder, 10 g/l Bacto tryptone, 5 g/l NaCl, 5 g/l sucrose crystallized, 5 g/l MgSO4·7H_2_O) and PSB medium (10 g/l peptone, 10 g/L sucrose crystallized, 1 g/l L(+)-glutamic acid, pH 7.4) were used for the OMVs release of *Xoc* BLS256. Antibiotics were added to the media in the following final concentrations (μg/ml): ampicillin, 100; rifampicin, 25; kanamycin, 25; and spectinomycin, 50.

### Outer membrane vesicles preparation

OMVs were isolated as previously described [22]. Briefly, make *Xoc* BLS256 starters (2 sterile tubes with 3 ml of sterile YEB medium in each), scrub 3-5 *Xoc* colonies from the plate, and add them to the medium in the tube. Put the tubes in 28 °C shaker for overnight. Then *Xoc* BLS256 starters were grown at 28°C in 500 ml PSB medium until OD_600_=0.6 for OMVs isolation. The cells were removed by low-speed centrifugation (8,000g) at 4°C, and the culture supernatant was filtered through a 0.22-μm-pore-size PVDF membrane filter (Millipore). OMVs were then collected from the filtered supernatant by centrifugation for 2 h in a Beckman Coulter Optima XPN-100 ultracentrifuge at 180,000g and 4°C. The OMV pellet was washed with OMV buffer (20 mM HEPES, 500 mM NaCl, pH 7.4) and re-pelleted by centrifugation at 200,000 g for 2 h at 4°C.

For OMVs purification, OMV pellets were re-suspended in OMV buffer in 60% OptiPrep Density Gradient Medium (Sigma-Aldrich, Germany) and layered with 0.8 ml 40% OptiPrep, 0.8 ml 35% OptiPrep, 1.6 ml 30% OptiPrep and 0.8 ml 20% OptiPrep. Samples were centrifuged for 16 h at 100,000 g and 4°C. 500 μl fractions were removed from the top of the gradient, with OMVs residing in fractions 2 and 3, corresponding to 25% OptiPrep, as previously shown [22]. The resulting pellets were resuspended in distilled water and filter-sterilized through 0.22-μm-pore-size PVDF membrane filter (Millipore) before beening stored at −80°C.

### Nanoparticle tracking analysis (NTA)

Nanoparticles in the isolated OMV suspensions were analyzed using a ZetaView PMX120 (Metrix) instrument. The platform was cleaned with 10 ml distilled water for 3 times to ensure no residue was present and to remove any potential air bubbles before analyzing samples. The cell with 100 nm alignment suspension was filled with platform by sterile syringes for standardization and then cleaned with 10 ml distilled water for 3 times. OMV suspensions were diluted in distilled water to a final volume of 600 μl and injected into the sample chamber with sterile syringes, yielding particle concentrations of 10^6^ particles per milliliter in accordance with the manufacturer’s recommendations. The software used for capturing and analyzing the data conformed the standard ASTM E2834-12.

### Transmission electron microscopy (TEM) imaging

Isolated OMVs sample were dropped onto copper grids for 2 min, and treated with 3% (w/v) phosphotungstic acid for 30 s. The samples were analyzed by Talos F200 transmission electron microscope (Thermo Fisher Scientific, USA) at 120 kV.

For TEM imaging, samples of the rice leaves were placed in 2.5% glutaraldehyde solution for at least 6 h, post fixed in osmium tetroxide, dehydrated in a graded series of ethanol, and embedded in LR White embedding media. Samples were polymerized at 60°C overnight and sectioned using a Leica EM UC7 microtome. The tissues of plant leaves were cut at 60-90 nm. TEM images were acquired using a Tecnai G2 spirit Biotwin microscope operated at 120 kV.

### RNA-Seq analysis of BLS256 sRNAs

Purified OMVs from BLS256 lysed with Qiazol reagent. OMV RNA was isolated with the miRNeasy kit (Qiagen), which retains the small RNA fraction. The RNA concentration was then determined with a Bioanalyzer (Agilent Technologies, Santa Clara, CA, USA). 1 μg total RNA was used for preparation of cDNA libraries with the TruSeq Small RNA Library Preparation Kit (Illumina, San Diego, CA, USA). Libraries were sequenced as 50 bp paired-end reads on an Illumina Genome Analyzer. Reads were trimmed and aligned to the BLS256 reference genome (NC_017267) using CLC Genomics Workbench (CLC-Bio/Qiagen). Data are deposited in NCBI under BioProject number PRJNA915483.

### Construction of deletion and complementary mutants

ΔXosr001 mutant strain was generated as described by Baba with minor modifications [23]. Two fragments flanking the target gene were amplified from the chromosomal DNA of *Xoc* BLS256 using Pfu polymerase (TransGen Biotech, Beijing, China) and the primers described in Table S2. The PCR products were digested, subcloned into the suicide vector pKMS1, and introduced into bacteria by electroporation (Bio-Rad Laboratories Inc., Hercules, CA, USA) with kanamycin selection. A single transformant with kanamycin resistance was selected, cultured for 8 h in NB, and inoculated as 10-fold dilutions to NA with 15% sucrose to obtain sucrose-insensitive clones.

To obtain the Xosr001 complementary mutant (ΔXosr001-pXosr), the full-length corresponding gene was amplified, and the fragment was cloned into pHM1 with the lac promoter. The recombinant plasmid was transferred into ΔXosr001 by electroporation, and transformants were screened on NA plates supplemented with spectinomycin.

### RNA sequencing and analysis of rice leaves

Total RNA was extracted from plant tissue at the site of inoculation using a MiniBEST Plant RNA Extraction Kit (TaKaRa, China) as per kit instructions. RNA was collected for two biological replicates for each condition. The concentration of total RNA was measured by the NanoDrop 3300 (Thermo Scientific, USA). mRNA was enriched from 4 μg total RNA by using magnetic beads linked with oligo dT. The obtained mRNA was fragmented into short fragments and used for first-strand cDNA synthesis and second strand cDNA synthesis. The resulting double-strands cDNA was purified by AMPure XP beads. Purified double-strands cDNA were then used for end repair, 3’ end adenosine tailing, sequencing adaptor connection, fragment selection and finally PCR amplification to generate sequencing library. The cDNA clusters were sequenced on the Illumina HiSeq 4000 platform using the paired-end technology and sequencing by synthesis method. The RNA-Seq data was submitted to NCBI database (https://www.ncbi.nlm.nih.gov/sra/) with SRA accession number PRJNA915483.

### Virulence assays

Virulence assays were conducted by spray and pressure inoculation [20, 24]. For pressure inoculation, *Xoc* strains were adjusted to an OD_600_ of 0.8 using needleless syringes. Lesion lengths were measured 14 d after inoculation. Twelve or more leaves were inoculated and evaluated for each *Xoc* strain. For spray inoculation, *Xoc* strains were re-suspended with 10 mM MgCl_2_ with 0.01% Silwet L77 (OD_600_=0.8), which were sprayed onto four-week-old seedlings of rice cv. Nipponbare. After 4 days post-inoculation (dpi), 3 pieces of 5 cm-long leaves were detached from the inoculated seedlings and ground in sterile water after surface sterilization with 75% ethanol with 3 technical repeats. The samples were spread on NA plates after serial dilution. Colony-forming units were counted after 2-day culturing.

### Stomatal conductance measurement

Stomatal conductance was measured using a photosynthesis system (Model LI-6400XT, Li-Cor Inc., Lincoln, NE, USA) [20]. Briefly, stomatal conductance was measured with a 2×3 cm leaf chamber supplied with a red-blue LED light source, and parameters were set as 400 μmol mol^-1^ CO_2_ and 200 μmol m^-2^ s^-1^ photosynthetic photon flux density (PPFD).

### Transgenic constructs and plant transformation

*OsJMT1* was amplified and cloned into pCAMBIA1300-35S::HA after digestion with *Eco*R I and *Hind*III. To construct a native promoter-driven expression vector, the full-length *OsJMT1* gene (~2 kb) was amplified from rice genomic DNA. The construct was transformed into *A. tumefaciens* strain EHA105 through the freeze-thaw method [25]. Transgenic rice plants were generated through *Agrobacterium*-mediated transformation of rice calli (cv. Nipponbare). The regenerated shoots were transferred into 1/2×MS medium for rooting and were then planted in the greenhouse.

Nippobare rice was genetically modified with CRISPR/Cas9 technology as described previously [26]. In brief, target sequences were selected the regions of *OsJMT1* (Table S1), and sgRNA was designed with CRISPR MultiTargeter (http://www.multicrispr.net/index.html) and synthesized by Shanghai Bioegene (Shanghai, China). The sgRNA and Cas9 constructs were transferred into Nippobare callus by Agrobacterium-mediated transformation (Biorun, Wuhan, China).

### RNA fluorescence in situ hybridization (RNA-FISH)

Samples of the rice leaves were placed in 50% FAA Fixative solution (Sigma-Aldrich, Germany) for at least 24 h at 4°C, re-hydrated by going through an ethanol series of 100, 85, and 75% respectively for twice (vol/vol) (5 min each) and DEPC solution (1min) at room temperature, and embedded in LR White embedding media. Samples were polymerized at 62°C for 2 h and sectioned using a Leica EM UC7 microtome. After protease (Sigma-Aldrich, Germany) digestion (20 min, 37°C), samples were treated with 0.2% glycine (Sigma-Aldrich, Germany) for 2 min, followed by a TEA treatment (Sigma-Aldrich, Germany) HCl and acetic anhydride (Sigma-Aldrich, Germany). After two washes in 1xPBS buffer, samples were de-hydrated and then hybridized with probes overnight at 53.3°C. For symbiont Xosr001 transcript and *OsJMT1* detection), RNA-FISH probes (Table S2) were designed and provided by Molecular Instruments (Sangon Biotech, China). Images were adjusted to optimize visual resolution using Leica Application Suite X (v. 3.4.2.18368).

### MeJA extraction and quantification

Endogenous MeJA was extracted from 4-week-old rice leaves. Briefly, the rice leaves (0.1 g) were ground to powder in liquid nitrogen. The powder was incubated overnight with 1 ml of PBS buffer (NaCl 8 g/L, KCl 0.2 g/L, Na_2_HPO_4_ 3.63 g/L, KH_2_PO_4_ 0.24 g/L, pH 7.4) at 4°C. The MeJA contents were quantified using plant MethylJasmonate (MeJA) ELISA Kit (Mlbio BioTech, China) following the manufacturer’s instructions.

### Measurement of sRNA expression level by semiquantitative RT-PCR

Total RNA of BLS256 pellets or BLS256 OMVs was isolated using the SurePrep™ RNA Purification Kit (Thermo Fisher Scientific). The cDNAs were synthesized from the RNA by reverse transcription using a RevertAid First Strand cDNA Synthesis Kit (Thermo Fisher Scientific) and used as the template for PCR amplification of a target sRNA gene. PCR was performed with a cycler using the following cycle parameters: 34 cycles of 94 °C for 15 s, 60 °C for 15 s, and 72 °C for 15 s. The amplification products were analyzed in 1.5% agarose gels and quantified using GelQuant.NET software. The 16S rRNA gene of *Xoc* BLS256 was used as the internal control to verify the absence of significant variation at the cDNA level in the two RNA samples.

### Western blot

To extract total proteins, 0.1 g rice leaf samples were chilled with liquid nitrogen and ground into a fine powder in a mortar, then homogenized in 700 μL of extraction buffer (125 mM Tris-HCl pH 6.8, containing 5% (w/v) SDS, 10% (v/v) Glycerol, and 2% (v/v) 2-mercaptoethanol). Total proteins were separated by 12% sodium dodecyl sulfate polyacrylamide gel electrophoresis (SDS-PAGE) and then transferred to polyvinylidene fluoride membranes (Merck Millipore, USA). OsJMT1-HA was detected by immunoblotting with HRP-conjugated anti-HA monoclonal antibodies (1:2,000, Roche), and the detection of OsActin using anti-Actin monoclonal antibody (1:5,000, CWBIO) was used as a loading control. Signals were visualized using Western Lightning Plus-ECL (Edo Biotech) and detected with ChemiScope 3000 mini (CLiNX, China).

### Northern blot

Total RNA was purified from *Xoc* strains liquid cultures (OD_600_ = 1.0) using the EasyPure RNA Kit (Transgen Biotech, China). Total RNA was extracted from plant leaves using a MiniBEST Plant RNA Extraction Kit (TaKaRa, China). RNA (10–20 μg) was separated in 1% agarose gels containing 25 mM guanidium thiocyanate, transferred to Hybond N+ nitrocellulose membranes (Merck Millipore, USA), and cross-linked to membranes by UV radiation. Probes were 5’-labeled with digoxygenin (DIG). Membranes were prehybridized for 10 min at 42°C, and then incubated with labeled probes overnight. Membranes were then rinsed, dried and visualized by phosphorimaging on a ChemiScope 3000 mini (CLiNX, China) as described previously [24].

### Quantitative real-time PCR

qRT-PCR was conducted as described previously [24]. Gene expression was normalized relative to *rpoD* or *OsActin* using the ΔΔCT method, where CT is the threshold cycle. Three independent biological replicates were included and analyzed using the Wilcoxon-Mann-Whitney test.

## Results

### OMVs contain differentially packaged sRNAs

The *Xoc* BLS256 OMVs were isolated using ultracentrifugation and purified by OptiPrep gradient density centrifugation. Analysis by TEM revealed the presence of distinct spheroid particles with an envelope structure and electron-dense luminal contents characteristic for OMVs (Figure 1a). The OMVs released by BLS256 ranged in size from 20 to 400 nm described by nanoparticle tracking analysis, with an average diameter of 120 nm as illustrated in Figure 1b. These results verified that we successfully extracted the *Xoc* BLS256 OMVs. RNA-Seq analysis of BLS256 OMVs was conducted to determine if sRNAs are packaged in OMVs, which revealed that the majority being ribosomal RNAs (58.03%) and tRNAs (21.79%), with a less complement of small ncRNAs (2.16%) (Figure S1). Specially, 3,615 unique sRNA sequences in OMVs was detected from RNA-Seq analysis. We listed 8 sRNAs with the highest predicted abundance in BLS256 OMVs and selected 4 sRNAs among them for validation (Table S3). The 8 candidate sRNAs were named Xosr001-Xosr008 *(Xanthomonas* OMV small RNA), with the ‘sr’ indicating that it is small RNA. Four intergenic sRNAs (Xosr001, Xosr002, Xosr003, Xosr006) identified by cDNA cloning method from OMVs (Figure 1c). Moreover, transcript lengths determined by northern blot analysis with sRNAs lengths confirmed the 4 sRNA candidates that we tested before (Figure 1d).

**Figure 1.**
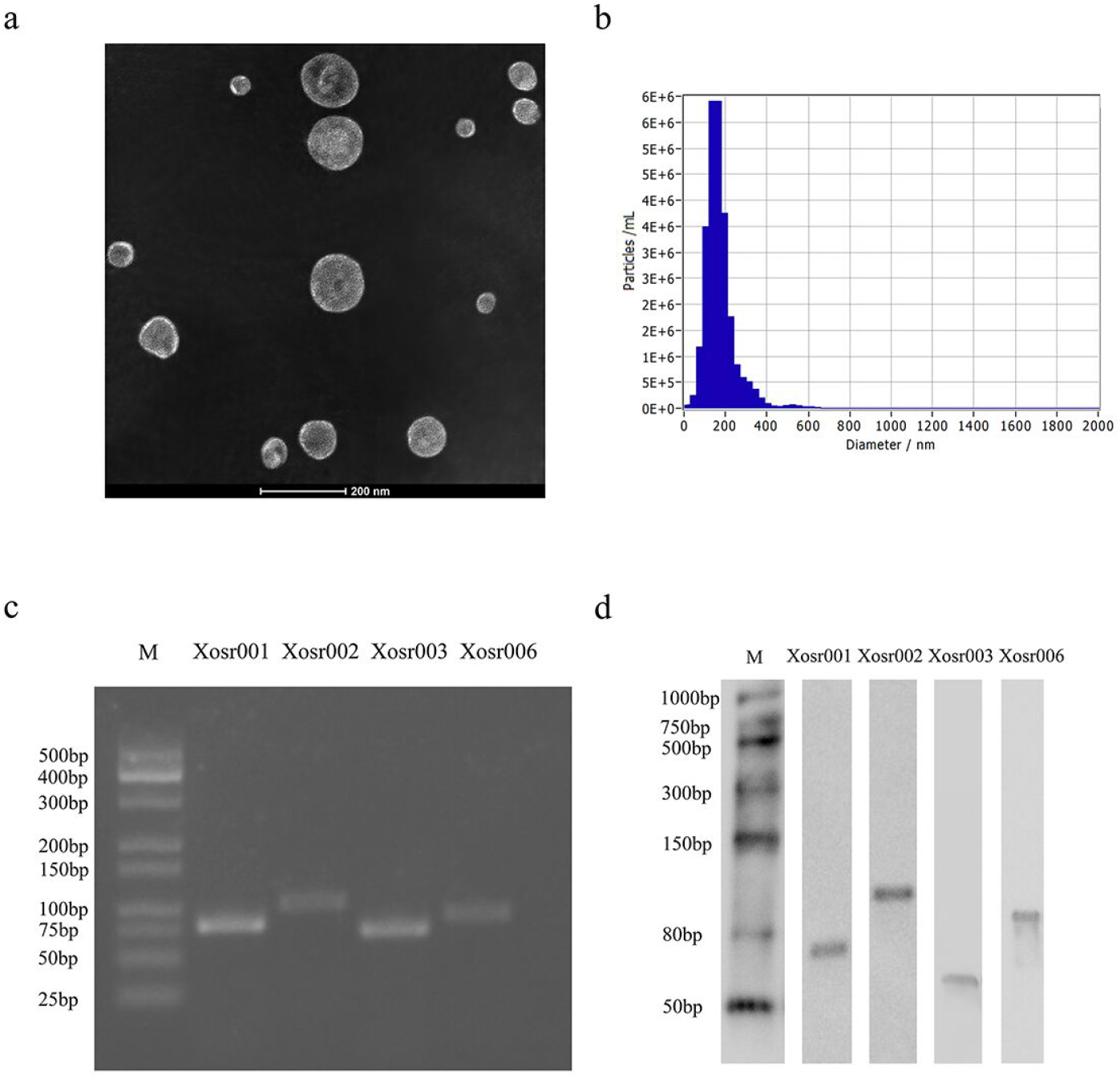
*Xoc* BLS256 releases OMVs enriched in sRNAs. (a) Transmission electron microscope (TEM) of OMVs purified from *Xoc* BLS256 following density step-gradient centrifugation. Size bar is 200 nm. (b) The size distribution of OMVs detected by nanoparticle tracking analysis (NTA). (c) PCR products of cDNA obtained in RT-PCR and fractionated on 3% agarose gel electrophoresis. Total RNA was isolated from purified BLS256 OMVs using TRIzol methods and Magic 1st cDNA Synthesis Kit (Magic-Bio, China) was used to generate cDNA. (d) Northern blots verify the presence of BLS256 OMVs sRNAs. Transcript sizes are approximate and compared with DIG-labeled marker (M).

### Xosr001 is transferred from OMVs to host plant cells

Tang et al. previously reported that sRNA designated sRX061 plays an important role in the virulence of *Xcc* [27]. Xosr001, the homologue of sRX061 in *Xoc* BLS256, was detected to be packaged in OMVs (Table S3). The secondary structure of Xosr001 was predicted using software available at https://sfold.wadsworth.org (Figure S2). The minority of reads mapped to sRNAs in OMVs by Illumina sequencing (Figure S1), which were found to have full coverage and, as such, appear not to be degraded. Xosr001 was the most abundant sRNA among them (Table S3). In addition, rRNAs found within OMVs, such as 16S rRNA, were also observed from RNA sequencing. The semi-quantitative RT-PCR analysis indicated that the expression level of Xosr001 in the bacterial cell compartment was roughly consistent with that in the purified OMVs. (Figure 2a and 2a’), suggesting there is no significant selective packaging of Xosr001 within OMVs.

**Figure 2.**
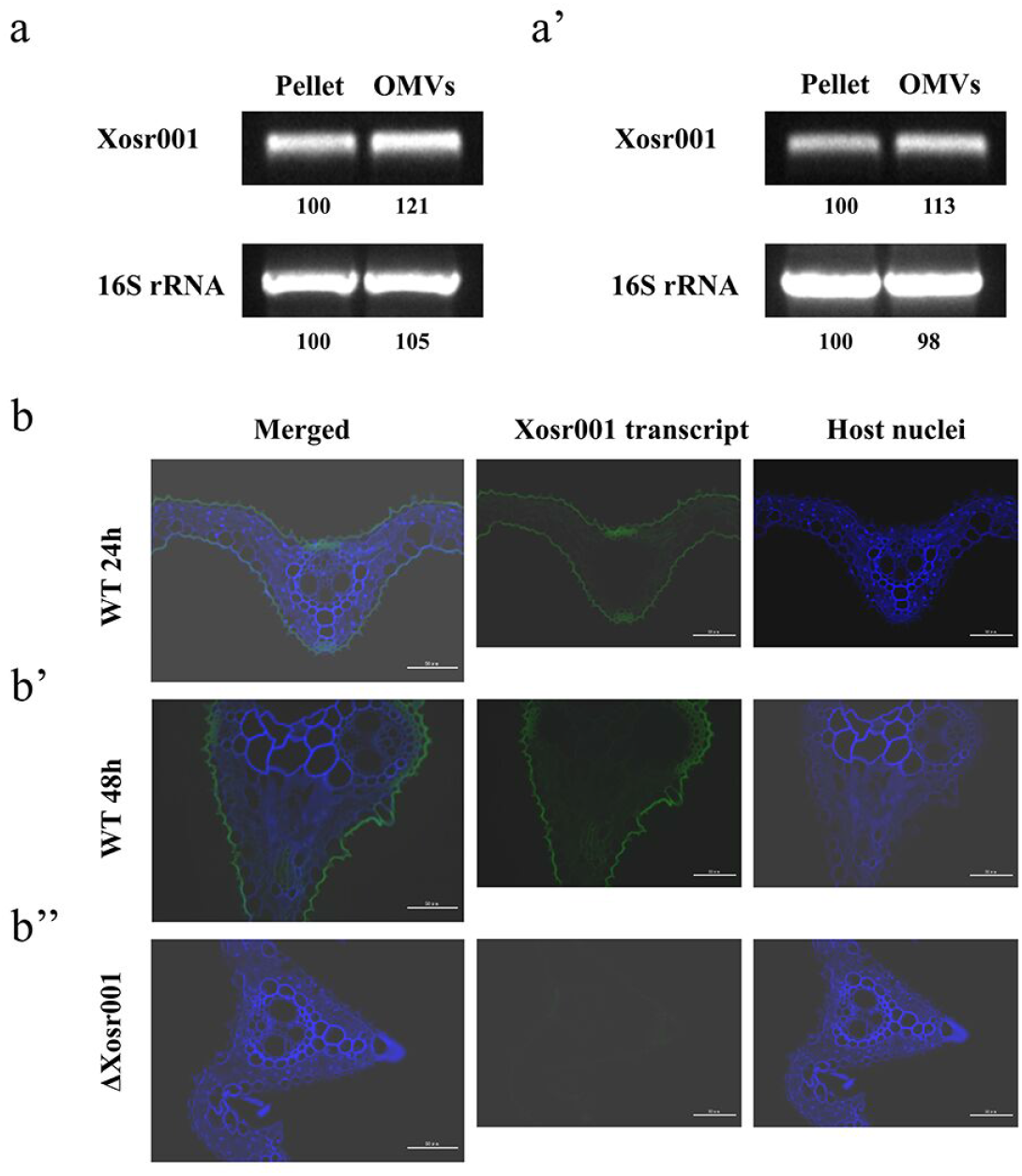
The noncoding sRNA, Xosr001, localizes within the epidermis and mechanical tissue of rice leaves. Detection of the expression level of Xosr001 in the BLS256 pellet and purified OMVs by semi-quantitative RT-PCR for 24 h (a) and 48 h (a’) cultivation respectively. Localization of Xosr001 transcript by confocal microscopy, 24 h (b) and 48 h (b’) after colonization by WT or ΔXosr001 (b”) bacteria. Left: merged images with orthogonal views; other panels: images of individual labels.

To determine whether Xosr001 enters into host plant tissues via OMVs, we contrasted its absence in the ΔXosr001 mutant to its localization in rice leaves colonized by WT inoculated for 24 h or 48 h. Using RNA fluorescence in situ hybridization (RNA-FISH), Xosr001 transcripts were found in WT-inoculated leaves (Figure 2b and 2b’) whereas no signal was detected in ΔXosr001-colonized ones (Figure 2b”). Significantly, the Xosr001 transcript signal was observed to be located in epidermis and mechanical tissue of rice leaves (Figure 2). Given the absence of evidence of bacterial lysis within the tissue of rice leaves by TEM (Figure S3c) and the biomass of bacteria have no significant differences after inoculation for 24 h or 48 h (Figure S3d), we hypothesized that the major source of the Xosr001 found within host plant cells is OMV-delivered using RNA-FISH. We conclude that the epidermis and mechanical tissue of rice leaves may be particularly susceptible to bacterial cargo, such as Xosr001 of *Xoc* BLS256, delivered by symbiont derived OMVs and secreted molecules.

### OMV-packaged Xosr001 trigger host plant response

The specific accumulation of plant defense gene transcripts is a key early component resulting in defense responses in infected tissue, which act as an elicitation signal for intercellular transmission in response to infection [28]. To investigate whether plant defense is influenced by the presence of Xosr001, we performed a comparative RNA-seq analysis on WT- and ΔXosr001-inoculated rice leaves 24 h after inoculation. Compared to WT-inoculated leaves, several host genes were significantly up-regulated in the ΔXosr001-inoculated rice leaves, including genes associated with the phytohormone synthesis pathway and stress-responsive genes with known immune-functions (Figure 3a).

**Figure 3.**
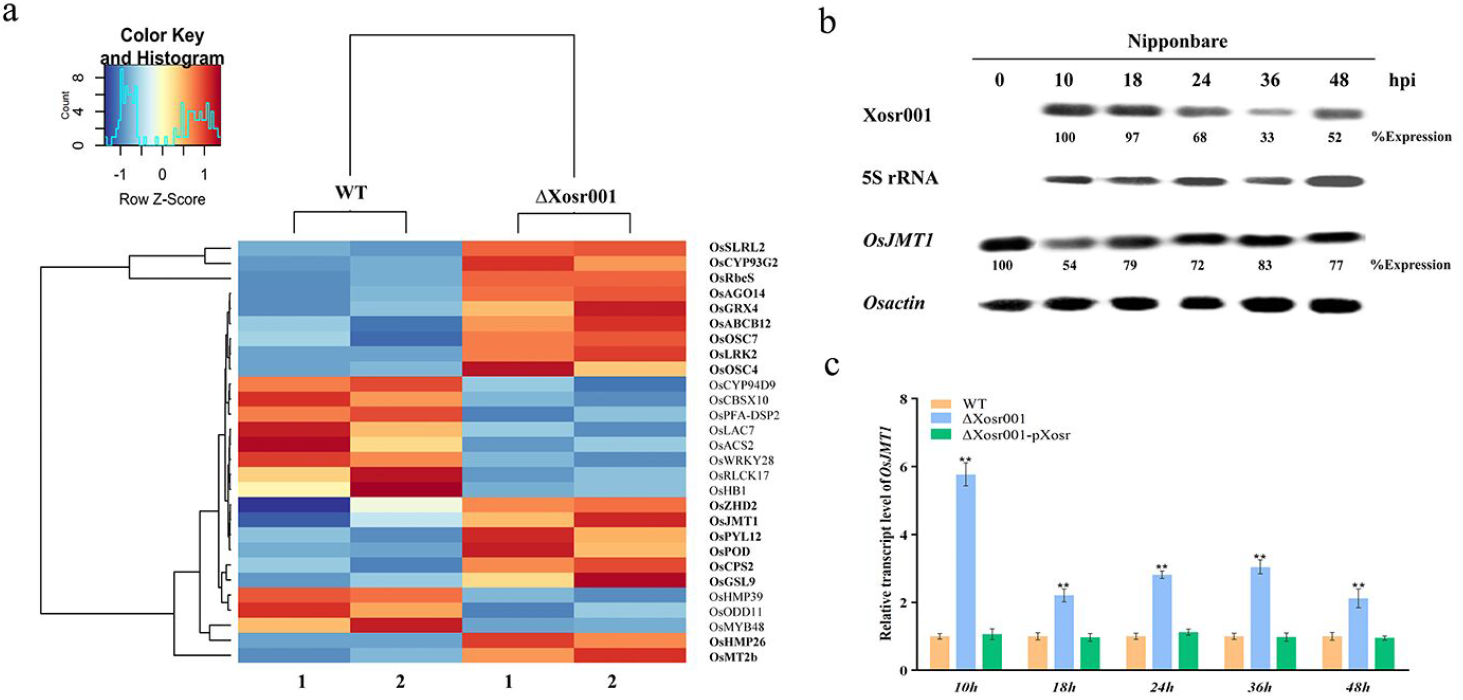
Host responses to colonization by WT or ΔXosr001 differ. (a) Heat map depicting fold-change differences in significantly differently expressed genes in leaves of rice cv. Nipponbare colonized by WT and the ΔXosr001 mutant. Genes that are up-regulated in ΔssrA-colonized leaves compared to WT-colonized are indicated in bold. The replicate number for each condition is indicated beneath the heat map. (b) Northern blot analysis of *Xosr001* and *OsJMT1* transcription levels were determined by inoculating six-week-old rice leaves with *Xoc* BLS256, respectively, for 0, 10, 18, 24, 36, 48 hours post-inoculation (hpi). 5S rRNA and *Osactin* were used as a loading control. Values above each band represent band intensity and were calculated using Image J software. Band intensity in the first lane was normalized as 100. (c) Expression levels of OsJMT1 after inoculating by WT, ΔXosr001, and ΔXosr001-pXosr for 0, 10, 18, 24, 36, 48 hpi respectively. Gene expression was detected by qRT-PCR using *OsActin* as an internal reference gene. Data are presented as means ± SE (n = 3 replicates per measurement). Asterisk (*) indicates a statistically significant difference in relative gene expression (two-sided t-test; *, P < 0.05; **, P < 0.01; ***, P < 0.001).

We were particularly interested in *OsJMT1*, which encodes a jasmonate methyl transferase, a key component of the plant immune response to pathogens, and converts JA to methyl jasmonate (MeJA) [29]. RNA-seq data indicated that the expression of *OsJMT1* was upregulated in the ΔXosr001 mutant as compared to the wild-type BLS256 (Figure 3a). To evaluate the expression of *OsJMT1* after inoculation, leaves of six-week-old rice cv. Nipponbare were inoculated with the WT for 10 h, 18 h, 24 h, 36 h, and 48 h, respectively. At 10 h post-inoculation (hpi), expression of *OsJMT1* was significantly reduced compared with non-inoculated leaves (Figure 3b, lanes 1 and 2). Interestingly, there was a substantial decrease in Xosr001 transcription at 36 hpi and the expression of *OsJMT1* was obvious increased when compared with 10 hpi (Figure 3b, lanes 2 and 5), which suggests that Xosr001 may promote *OsJMT1* transcripts degradation. RT-qPCR results correlated with the northern blot analyses and showed elevated expression of *OsJMT1* transcripts after 10 hpi (Figure 3c). Collectively, these results suggest that the presence of OMV-packaged Xosr001 sensing generates a dysregulated host plant response.

### Xosr001 promotes OsJMT1 degradation and endogenous MeJA content

To further evaluate whether *OsJMT1* transcript was associated with Xosr001 *in vivo* and where it occurred within the plant tissues. RNA-FISH harboring specific RNA probes of Xosr001 and *OsJMT1* was performed to determine that Xosr001 suppressed OsJMT1 expression *in vivo*. Leaves of six-week-old rice cv. Nipponbare were inoculated by WT and ΔXosr001, while WT-inoculated leaves served as a negative control. As expected, mutant ΔXosr001-inoculated leaves showed enhanced *OsJMT1* transcription when compared to WT (Figure 4a and a’) at 24 hpi. Moreover, the luminescence of *OsJMT1* transcripts in WT-inoculated leaves emitted approximately 40% of the ΔXosr001 level (Figure 4b), suggesting that BLS256 OMVs lacking Xosr001 strengthened *OsJMT1* expressed in rice leaves. In addition, *OsJMT1* transcripts were observed not only inside the epidermis but also within the mechanical tissue of rice leaves and directly contacted the OMV-packaged Xosr001 (Figure 4a), which indicated that Xosr001 reduced the *OsJMT1* transcription level only when it was transferred into the epidermis and mechanical tissue of rice leaves.

**Figure 4.**
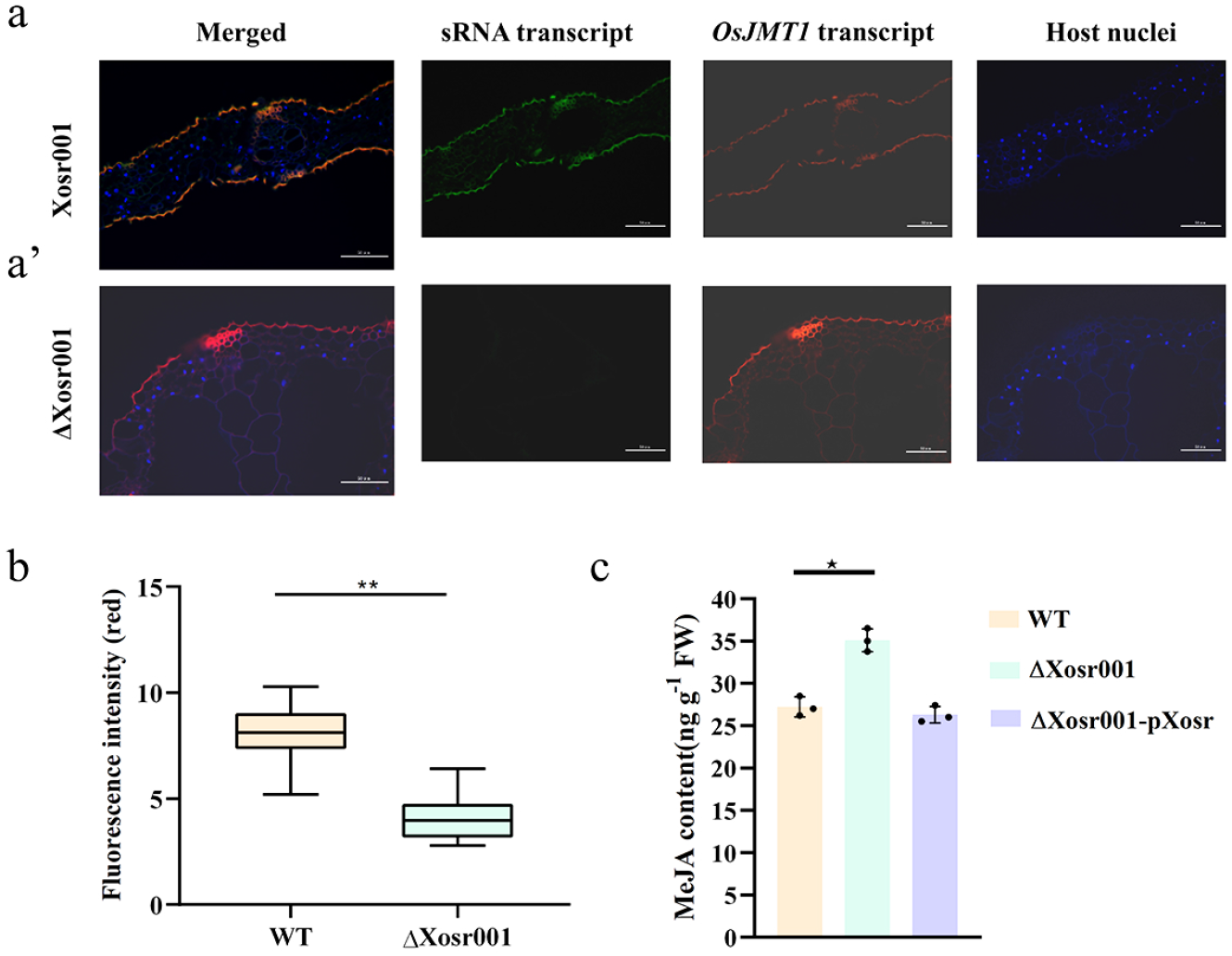
Localization of the *OsJMT1* transcript (magenta) and Xosr001 transcript (green) using hybridization chain-reaction fluorescence in situ hybridization labeling. Leaves of six-week-old rice cv. Nipponbare were inoculated by WT (a) and ΔXosr001 (a’) at 24 hpi. (b) Quantification of laccase-3 signal using relative fluorescence intensity of a Z-series image of the light organ (n = 5). P values were calculated using a 1-way ANOVA with TMC (**, P < 0.01). (c) Endogenous MeJA level of rice leaves. Leaves of six-week-old rice cv. Nipponbare were inoculated by WT, ΔXosr001, and ΔXosr001-pXosr respectively for 24 hpi. All data are presented as the mean±s.e.m. Asterisk (*) indicate significant differences at P<0.05 (ANOVA and Duncan’s multiple range test), n=3.

Previous studies have demonstrated that OsJMT1 converts JA to MeJA [29]. To assess whether Xosr001 has an impact on the contents of endogenous MeJA, we generated a BLS256 deletion mutant for Xosr001 as well as a re-complemented mutant that stably expressed Xosr001 from a plasmid with a lac promoter (here after called ΔXosr001-pXosr). ΔXosr001 showed significantly enhanced endogenous MeJA contents in rice leaves at 24 hpi as compared to the WT and there was no significant difference in the ΔXosr001-pXosr (Figure 4c). Thus, our results suggest that Xosr001 caused a reduction in *OsJMT1* transcript levels thereby inhibiting endogenous MeJA contents.

### Xosr001 induce stomatal immunity to enhance disease susceptibility in rice

In plants, MeJA could induce stomatal immunity to prevent phytopathogenic bacteria from entering host tissues during initial infection [19]. To investigate whether Xosr001 disarms stomatal immunity, stomatal conductance (Gs) was measured for rice leaves after spray inoculation. The water-treated rice seedlings had significantly higher Gs than regardless of *Xoc* strains and *E.coli* after 1 dpi (Figure 5a). Compared to WT and ΔXosr001-pXosr, there was a substantial decrease in Gs of ΔXosr001 at 2 dpi. Nevertheless, we observed that *E.coli*-inoculated rice leaves showed no significant difference in Gs after 1 and 2 dpi (Figure 5a). These results suggested that pathogenic bacteria *Xoc* strains could mediate stomatal opening, and the ΔXosr001 mutant obviously impaired the ability of stomatal re-opening on rice leaves, whereas nonpathogenic bacteria, such as *E.coli*, could not induce stomatal immunity in a nonhost plant. Collectively, Xosr001 suppresses *Xoc*-mediated stomatal opening on rice leaves. To evaluate whether Xosr001 contributes to disease susceptibility in rice, leaves of six-week-old rice cv. Nipponbare were inoculated by spraying with the WT, ΔXosr001 and ΔXosr001-pXosr. The ΔXosr001 mutant showed obviously degraded virulence and produced slightly shorter lesions than the WT and ΔXosr001-pXosr (Figure 5b and c). Consistent with disease symptoms, *in planta* bacterial population of *Xoc* BLS256 was significantly greater than that of ΔXosr001 (Figure 5d), indicating that Xosr001 contributed to the virulence in rice leaves through stomatal immunity.

**Figure 5.**
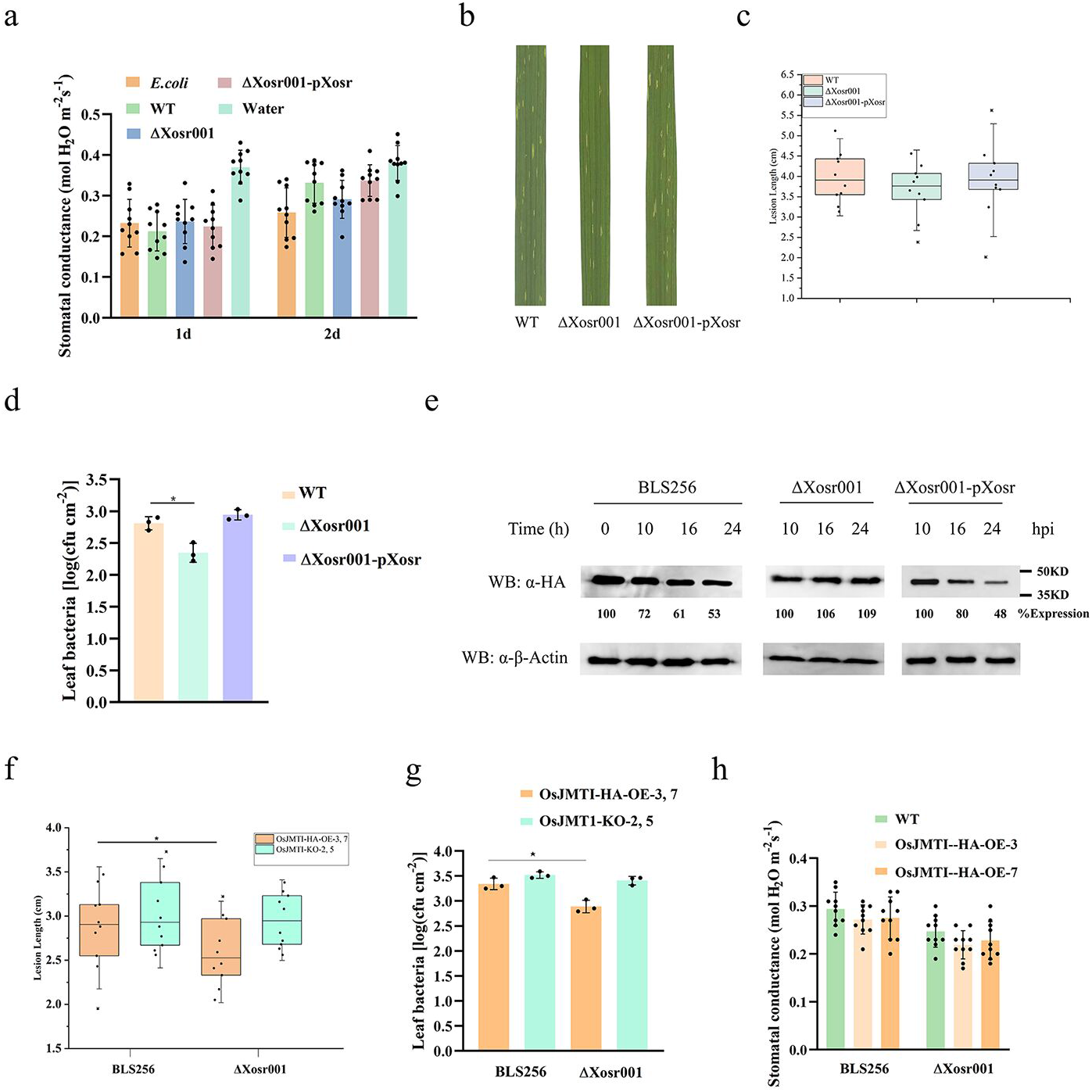
(a) Stomatal conductance in the leaves of 6-week-old seedlings of rice cv. Nipponbare by spray inoculation after challenging with WT, ΔXosr001, ΔXosr001-pXosr, *E.coli*, and water. Stomatal conductance was measured at 1 day post-inoculation (dpi) and 2 dpi. Data are shown as means ± SE (n = 10 technical replicates per measurement). Disease lesions (b) and lesion lengths (c) on the WT, ΔXosr001, and ΔXosr001-pXosr-inoculated rice leaves after spray inoculation. Six-week-old seedlings were sprayed with 0.01% Silwet L77. Photographs were taken at 4 dpi. Results indicate means ± SD. ANOVA was performed with Dunnett’s multiple comparison post-hoc correction as compared with the WT. (d) Bacterial population sizes in the WT, ΔXosr001, and ΔXosr001-pXosr-inoculated leaves Bacterial populations were determined at 4 days post-inoculation (dpi). (e) OsJMT1-HA was obviously degraded during *Xoc* infection, but remained relatively stable after ΔXosr001 infection. Three-week-old transgenic seedlings constitutively expressing OsJMT1-HA were sprayed with the strains BLS256, ΔXosr001, ΔXosr001-pXosr. OsJMT1-HA was detected by immunoblotting at the indicated time-points post inoculation. (f) Virulence was assessed by inoculating three-week-old susceptible rice plants. Leaves (n = 10) were inoculated with needleless syringes, and lesion lengths were evaluated 14 d after inoculation. Results indicate means ± SD. ANOVA was performed with Dunnett’s multiple comparison post-hoc correction as compared with the WT (*, P < 0.05). (g) Bacterial population sizes in the BLS256 and ΔXosr001-inoculated leaves of the OsJMT1-HA-OE and OsJMT1-KO transgenic plants. Bacterial population was determined at 4 dpi.Data are shown as means ± standard error (SE) (n = 3 technical replicates per measurement). The letters (d, g) indicate a statistically significant difference in bacterial population sizes as revealed by one-way ANOVA, Tukey’s honest significance test. (h) Stomatal conductance in the leaves of the 3-week-old OsJMT1-HA-OE transgenic lines (OsJMT1-HA-OE-3 and OsJMT1-HA-OE-7) after challenging with BLS256 and ΔXosr001 respectively. Stomatal conductance was measured at 2 dpi. Data are shown as means ± SE (n = 10 technical replicates per measurement).

We generated OsJMT1-HA-OE transgenic rice plants expressing OsJMT1-HA driven by the CaMV 35S promoters and obtained OsJMT1 knockout mutants OsJMT1-KO using CRISPR/Cas9-based gene editing to provide additional support for the observation that Xosr001 delivered into host plant cells by OMVs reduces OsJMT1 transcription and thus induces stomatal immunity. Expression of OsJMT1-HA in those transgenic lines was detected by immunoblotting (Figure S4). Spray inoculation of the WT and ΔXosr001-pXosr caused evident degradation of OsJMT1-HA in OsJMT1-HA-OE transgenic seedlings, while OsJMT1-HA was stable in the ΔXosr001-inoculated transgenic seedlings (Figure 5e). After spray inoculation with ΔXosr001, OsJMT1-HA-OE transgenic plants exhibited significantly shorter lesions and lower bacterial populations than those inoculated with WT, whereas ΔXosr001 or WT-inoculated leaves of OsJMT1-KO transgenic plants showed no obvious difference in virulence (Figure 5f and g). Besides, the absence of Xosr001, significantly compromised stomatal closure in OsJMT1-HA-OE transgenic plants after *Xoc* strains infection (Figure 5h). Thus, our findings indicate that Xosr001 influence the *OsJMT1* transcription level to induce stomatal re-opening during the infection process.

## Discussion

In this report we show for the first time that cross-kingdom regulation by sRNA contained in phytopathogenic bacterial OMVs. Our studies demonstrate that Xosr001 packaged in OMVs by *Xoc* BLS256 is a novel mechanism of immunomodulatory nucleic acids that drive host plant immune responses.

Recently several studies have described that microbial-derived RNA can act as a PAMP to facilitate infection via hijacking the RNA interference machinery of the host [6, 7, 30]. In light of these reports, vesicle-mediated could transport sRNAs into host tissue which are important players in gene expression regulation and aligned with specific regions of specific host transcripts. The periodontal pathogen *Porphyromonas gingivalis* driven sRNA23392, a 20-nt sRNA, can be conveyed by OMVs and targets the desmocollin-2 (DSC2) to promote the invasion and migration of oral squamous cell carcinoma (OSCC) cells [13]. Moreover, with the development of bioinformatics approaches, long sRNAs (30-100 nt) enriched in bacterial OMVs can be detected using high-throughput sequencing [12, 31]. In this study, we identified sRNA Xosr001 (77 nt) as being enriched in *Xoc* OMVs and capable of being delivered into host plant cells (Figure1 and 2), suggesting differential packaging of sRNAs is a significant portion of the OMV-associated RNA, although little known about how sRNAs are packaged into OMVs.

In plants, sRNAs from fungal plant pathogens can be delivered into the host plant to regulate endogenous plant genes, such as *Botrytis cinerea*, which produces sRNAs (Bc-sRNAs) that move into the host plant cell and hijack the plant RNA interference machinery [32]. This raises the possibility that OMVs-derived trans-kingdom gene silencing could occur between plants and phytopathogenic bacteria. In this report, we showed that sRNA Xosr001 has high abundance in the phytopathogenic bacterium *Xoc* BLS256 OMVs and RNA-FISH assays revealed that Xosr001 could decrease the expression of *OsJMT1* in rice leaves by blocking transcription processing (Figure 3 and 4), suggesting sRNA molecule can participate in cross-kingdom communication in *Xoc* and rice interactions. Although *Xoc* OMV-associated sRNA has not been reported previously, several groups have characterized the intracellular sRNA content of *Xanthomonas* using RNA-Seq and investigated that sRNA-mediated virulence regulation pathway under different growth conditions [24, 27, 33]. Tang et al. recently reported that particular sRNA, RsmU, acted as a negative regulator in the virulence and cell motility of *Xcc* [27]. Moreover, our previous study found that sRNA Xonc3711 of *Xoc* played an extensive role in oxidative stress and the biosynthesis of flagella by modulating the DNA-binding protein Xoc_3982 transcription [24]. Collectively, sRNAs could be helpful in understanding the virulence and biological circuitry of phytopathogenic *Xanthomonas* spp.

Disabling host stomatal immunity is usually a prerequisite for entering into leaf tissue successfully by various phytopathogenic bacteria [34]. Several derivatives of jasmonates (JAs) are found naturally in plants [35]. Based on the current reports that some of which are available, in particular MeJA, could close the stomatal pore [36, 37]. On the basis of our observations that OMVs-assisted delivery of *Xoc* sRNA Xosr001 is necessary for reduction of *OsJMT1* transcripts thereby influences the endogenous MeJA contents during the infection in rice (Figure 4). Furthermore, our data showed that significantly compromised stomatal closure was detected in ΔXosr001 when compared to *Xoc* BLS256 after 2 dpi (Figure 5a). More convincingly, the transgenic rice lines OsJMT1-HA-OE exhibited an attenuated stomatal immunity and disease susceptibility after ΔXosr001 infection compared with *Xoc* BLS256 infection (Figure 5f-h). These results indicate that *Xoc* OMVs-secreted Xosr001 enhances the stomatal immunity of rice during infection. However, the mechanism(s) by which Xosr001 directly target the *OsJMT1* transcripts remains to be determined, while the possibilities include sequence specificity or secondary structure playing a role in the recognition of target gene in host by Xosr001.

Interestingly, OsJMT1-HA was obviously degraded in *Xoc*-inoculated rice leaves at 24 hpi (Figure 5e), suggesting that Xosr001 promotes OsJMT1-HA degradation not only in stomata on the context of epidermal tissues but also in other tissue cells. This result is consistent with our observation that Xosr001 and *OsJMT1* transcripts were detected in the epidermis as well as in the mechanical tissue of rice leaves (Figure 4a and 4a’). Therefore, Xosr001 might have other potential functions in the mechanical tissue besides inhibition of stomatal immunity. It has been reported that sRX061, the homologue of Xosr001, regulated multiple systems including virulence, hypersensitive response, and swarming motility in *Xcc* [27]. It will be fully interesting to elucidate that Xosr001 also serves as a virulence effector using other mechanisms for control of bacterial infections in the future.

In summary, we uncover the role of OMV sRNA Xosr001 in pathogenic bacteria-host plant interactions. Xosr001 could be packaged in OMVs of *Xoc* BLS256 and loaded into the epidermis and mechanical tissue of rice leaves, which inhibit the transcription of *OsJMT1* to attenuate the endogenous MeJA contents thereby trigger the stomatal immunity in rice (Figure 6). Our findings highlight a novel approach to understand the cross-kingdom RNAi between phytopathogenic bacteria and host plants.

**Figure 6.**
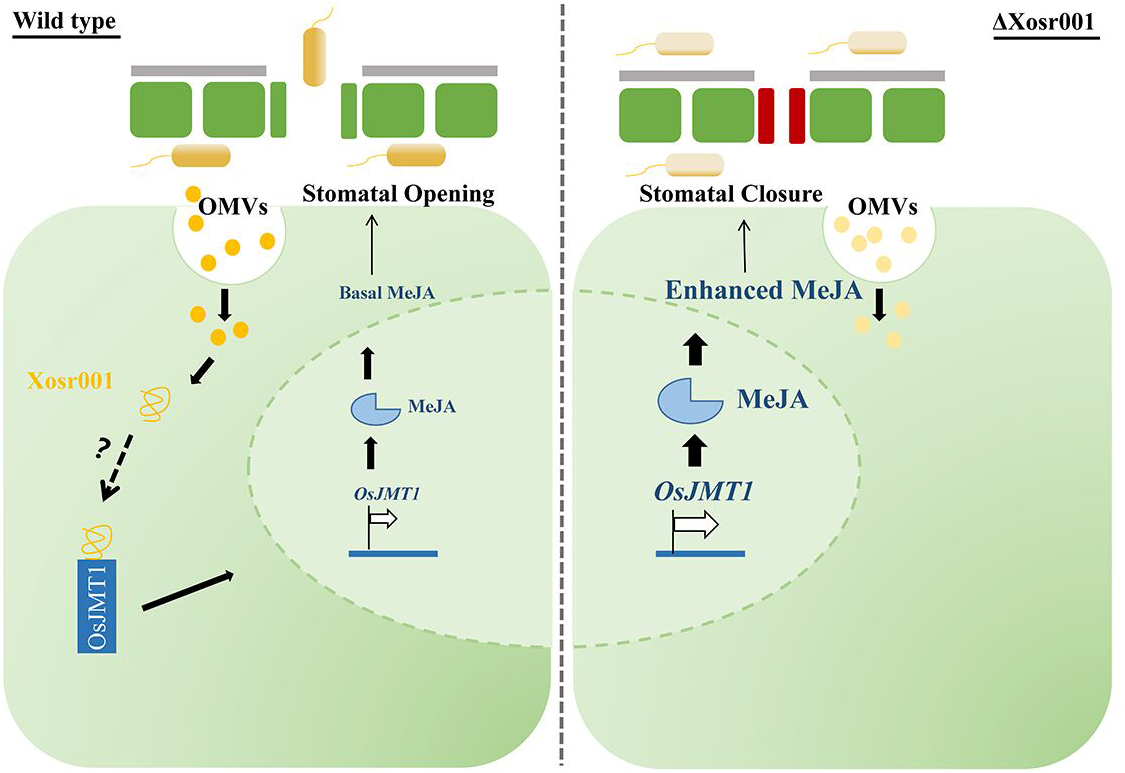
A working model for Xosr001 function in promoting MeJA contents and suppressing stomatal immunity. During *Xoc* infection, sRNA Xosr001 is secreted into host cells via OMVs, which inhibit the transcription of OsJMT1, and thus enhances endogenous MeJA contents and suppresses stomatal reopening thereby stomatal defense is attenuated for the successful entry of the pathogen.

## Data availability

The RNA-Seq data was submitted to NCBI database (https://www.ncbi.nlm.nih.gov/sra/) with SRA accession number PRJNA558244.

## Acknowledgements

This work was supported by National Natural Science Foundation of China (32272479, 32200142), Open Project Program of State Key Laboratory of Rice Biology (20190109), Open Project Program of State Key Laboratory for Biology of Plant Diseases and Insect Pests (SKLOF202201), Zhejiang Province ecological Environment Research and Promotion Project (2020HT0009), Shanghai Committee of Science and Technology (19390743300 and 21ZR1435500) and Chongqing Natural Science Foundation (CSTB2022NSCQ-MSX0524)

---

Figure S1 Relative proportions of types of *Xoc* BLS256 RNAs present in the RNA cargo of OMVs.

Figure S2 Predicted secondary structure of Xosr001. Predominant features in the secondary structure are labeled as follows: stems (S1–S5), bulges (B1-B2), loops (L1–L3), and single-stranded regions (SS1).

Figure S3 The biomass of bacteria have no significant differences after inoculation for 24 h (a) or 48 h (b). (c) Bacteria count were randomly measured from 5 parallel views by transmission electron microscopy. Wilcoxon rank-sum test. (d) Expression levels of specific gene *xoc_2071* of BLS256, which were calculated relative to *rpoD* using the ΔΔCT method, where CT is the threshold cycle. The 3-week-old seedlings were inoculated with *Xoc* BLS256 for 24 h or 48 h. Four independent biological replicates were carried out in this experiment.

Figure S4 The expression of OsJMT1-HA under the 35S promoter in the OE-3 and OE-7 transgenic rice lines was detected by immunoblotting.

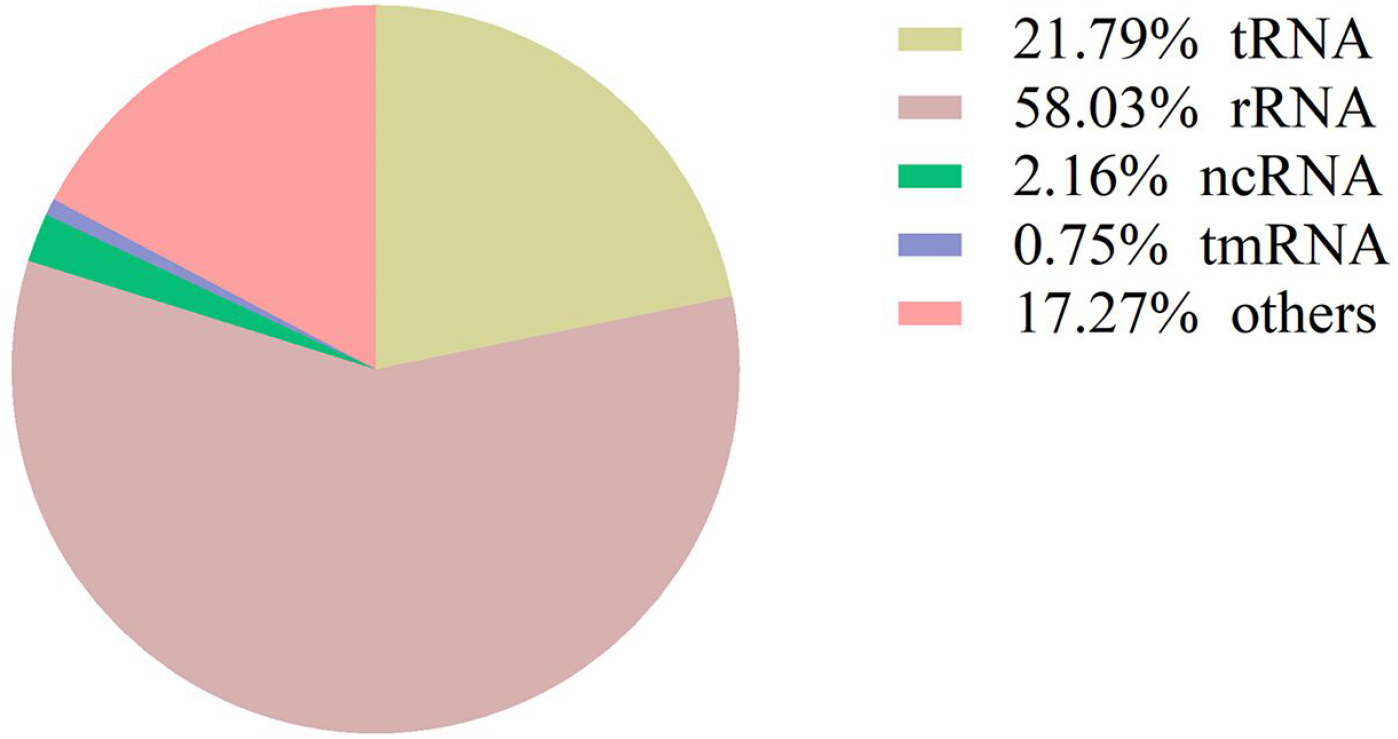

